# Loss of heterochromatin at endogenous retroviruses creates competition for transcription factor binding

**DOI:** 10.1101/2022.04.28.489907

**Authors:** Ryan O’Hara, Laura A. Banaszynski

**Affiliations:** Cecil H. and Ida Green Center for Reproductive Biology Sciences, Department of Obstetrics and Gynecology, Children’s Medical Center Research Institute, Harold. C. Simmons; Comprehensive Cancer Center, Hamon Center for Regenerative Science and Medicine, University of Texas Southwestern Medical Center, Dallas, Texas 75390, USA

## Abstract

The mammalian genome is partitioned into active and inactive regions, broadly termed euchromatin and heterochromatin, respectively. The majority of heterochromatin consists of repetitive elements, including endogenous retroviruses (ERVs). ERVs are enriched in regulatory elements containing transcription factor (TF) binding sites with individual families containing hundreds to thousands of distinct copies scattered throughout the genome. We hypothesized that epigenetic derepression of ERVs (such as that observed during early development) may alter the stoichiometry between TFs and their euchromatic target sites, with ERVs effectively competing for these factors. To test this, we modeled acute heterochromatin loss using inducible deletion of the co-repressor KAP1 in mouse embryonic stem cells (ESCs). Upon KAP1 deletion, we observe clear reductions in chromatin accessibility, histone acetylation, and TF binding at euchromatic regions. To directly test the concept of global binding site competition, we designed exogenous binding site arrays (EBSAs) to introduce upwards of 1500 copies of the OCT4 TF binding motif into ESCs. OCT4 EBSAs specifically reduce chromatin accessibility at POU family motifs and result in reduced transcription of the pluripotency machinery with subsequent differentiation. Overall, these data support a model in which heterochromatin at ERVs promotes euchromatic TF binding and transcriptional homoeostasis. We propose that regulated ERV derepression during pre-implantation may serve as a developmental siphon to weaken the robustness of ongoing transcription programs in favor of the plasticity required for cell fate specification.

## Main Text

The majority of heterochromatin in mammalian cells consists of repetitive elements, namely long and short interspersed elements (LINEs, SINEs) and endogenous retroviruses (ERVs) (*1*). In mouse embryonic stem cells (ESCs), heterochromatin at ERVs is maintained by the tripartite motif-containing 28 (TRIM28 also known as KAP1) co-repressor / SET Domain Bifurcated Histone Lysine Methyltransferase I (SETDB1 also known as ESET) complex that installs trimethylation on histone H3 at lysine 9 (H3K9me3) at ERVs (*2, 3*) (fig. S1). Loss of heterochromatin via acute deletion of either one of these factors leads to transcriptional derepression of specific ERV families accompanied by the accumulation of acetylation of lysine 27 on histone H3 (H3K27ac), a chromatin modification associated with active regulatory elements (*2*–*4*). Interestingly, the long terminal repeats (LTRs) of ERVs contain numerous sites for transcription factor binding and have been shown to act as enhancers for nearby genes (*4*–*9*). Of note, individual ERV families contain hundreds to thousands of distinct copies scattered throughout the genome (*10*), leading to the intriguing hypothesis that global derepression of TF binding sites within ERVs may alter the stoichiometry between TFs and their euchromatic target sites, effectively competing for these factors.

To test whether loss of KAP1 influences transcription factor binding at euchromatin, we used a conditional knockout mouse embryonic stem cell (ESC) line in which treatment with 4-hydroxytamoxifen (4-OHT) results in gene deletion and progressive loss of KAP1 protein over the course of 3 days (Fig. 1A). Loss of KAP1 does not appreciably affect expression of the Oct4 transcription factor over this time period, suggesting that these ESCs maintain pluripotency up to 75 hours after KAP1 deletion (Fig. 1A). Using chromatin immunoprecipitation followed by quantitative PCR (ChIP-qPCR), we found that KAP1 deletion resulted in loss of H3K9me3 enrichment at ERVs sequences such as intracisternal A-type particles IAPA and IAPEz (Fig. 1B). Loss of H3K9me3 preceded an increase in H3K27ac at these regions at later time points (Fig. 1C). These results are consistent with the literature and previously reported transcriptional activation of ERVs upon loss of heterochromatin (*2, 3, 11*). Interestingly, KAP1 deletion also resulted in reduced H3K27ac enrichment at annotated enhancers nearby the *Pou5f1* (Oct4) and *Klf4* genes as observed by ChIP-qPCR (Fig. 1D). We confirmed these results using H3K27ac Cleavage Under Targets and Tagmentation (CUT&Tag) in control and KAP1 deletion ESCs, revealing increased H3K27ac enrichment at several ERV subfamilies (e.g., IAPEz, IAPEY, MERVK9E) and decreased H3K27ac at regulatory elements of *Pou5f1* and *Klf4* (Fig. 1E, F). Reduced acetylation at euchromatin (defined as H3K27ac regions) upon loss of KAP1-mediated heterochromatin was also evident genome-wide (Fig. 1G,H).

**Figure 1.**
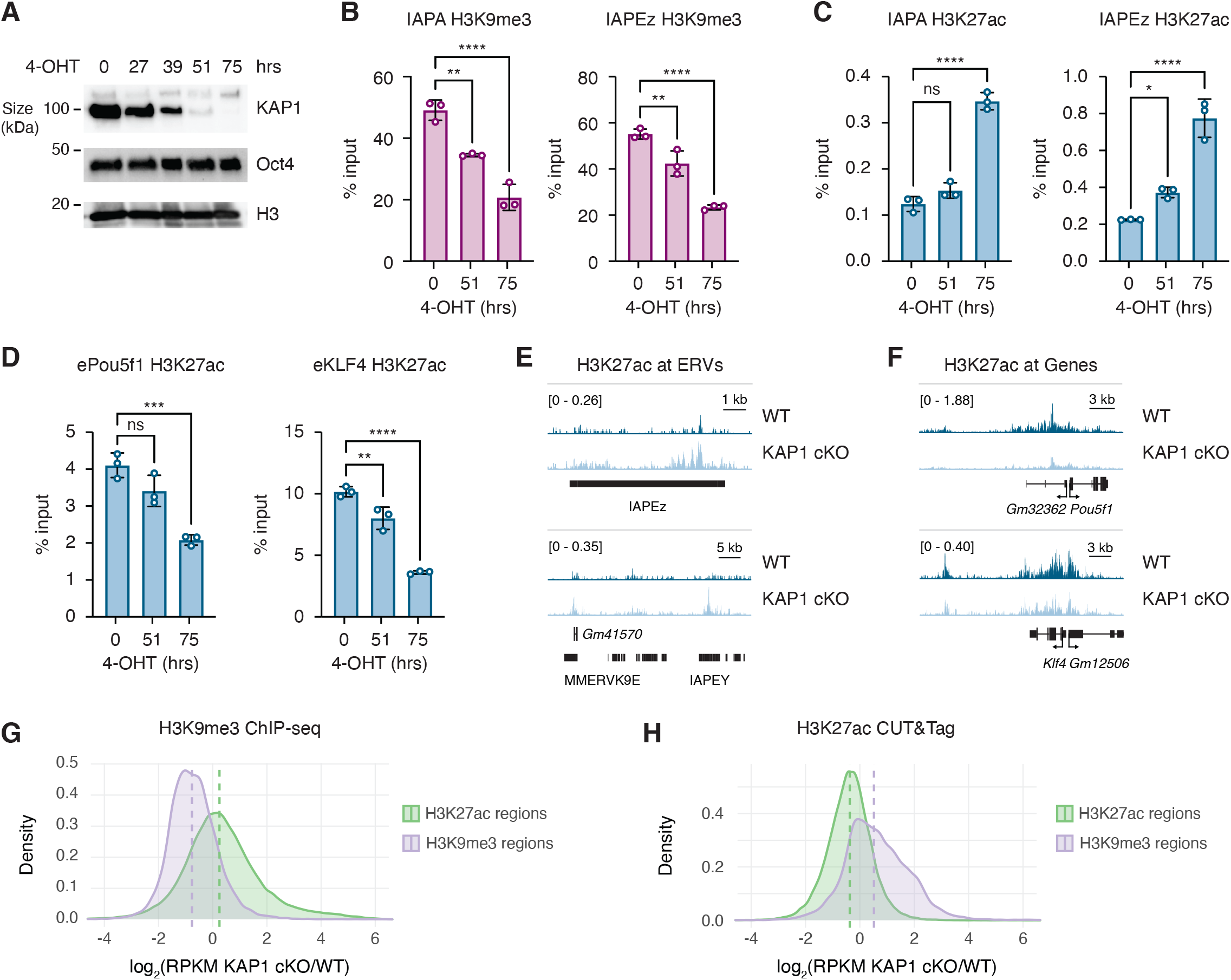
Loss of the KAP1 co-repressor reduces euchromatic histone acetylation. **(A)** Immunoblot from whole-cell lysates of KAP1 cKO ESCs treated with 1 μM 4-OHT for the indicated times. **(B)** ChIP-qPCR of H3K9me3 at IAPA and IAPEz ERVs in KAP1 cKO ESCs treated with 1 μM 4-OHT for the indicated times. Data are normalized to input and represent mean +/- s.d., ordinary one-way ANOVA, **p=0.0022 (0 vs 51 hr), ****p<0.0001 (0 vs 75 hr) for IAPA, **p=0.007 (0 vs 51 hr), ****p<0.0001 (0 vs 75 hr) for IAPEz. **(C)** ChIP-qPCR of H3K27ac at IAPA and IAPEz ERVs in KAP1 cKO ESCs treated with 1 μM 4-OHT for the indicated times. Data are normalized to input and represent mean +/- s.d., ordinary one-way ANOVA, ns=0.1406 (0 vs 51 hr), ****p<0.0001 (0 vs 75 hr) for IAPA, *p=0.0468 (0 vs 51 hr), ****p<0.0001 (0 vs 75 hr) for IAPEz. **(D)** ChIP-qPCR of H3K27ac at enhancers in KAP1 cKO ESCs treated with 1 μM 4-OHT for the indicated times. Data are normalized to input and represent mean +/- s.d., ordinary one-way ANOVA, ns=0.0662 (0 vs 51 hr), ***p=0.0005 (0 vs 75 hr) for ePou5f1, **p=0.0069 (0 vs 51 hr), ****p<0.0001 (0 vs 75 hr) for eKlf4. **(E)** Genome browser representations of H3K27ac ChIP-seq enrichment at ERVs in KAP1 cKO ESCs untreated (WT) or treated with 1 μM 4-OHT for 72 hrs (KAP1 cKO). The y axis represents read density in counts per million (CPM). **(F)** Genome browser representations of H3K27ac ChIP-seq enrichment at genic regions in KAP1 cKO ESCs untreated (WT) or treated with 1 μM 4-OHT for 72 hrs (KAP1 cKO). The y axis represents read density in counts per million (CPM). **(G)** Ratio (log2) of H3K9me3 CUT&Tag enrichment at H3K9me3-enriched and H3K27ac-enriched regions in WT and KAP1 cKO ESCs. **(H)** Ratio (log2) of H3K27ac CUT&Tag enrichment at H3K9me3-enriched and H3K27ac-enriched regions in WT and KAP1 cKO ESCs. For panels G and H, x axis values <0 indicate reduced enrichment in the absence of KAP1.

Histone acetylation, and particularly H3K27ac, broadly defines transcriptionally active regions of the genome. This modification is put in place by histone acetyltransferases that are recruited to target sites through the sequence-specific binding of transcription factors (TFs) to short motifs found at regulatory elements (*12*–*14*). The ability of TFs to bind target sites in part depends on chromatin accessibility, and methods such as assay for transposase-accessible chromatin using sequencing (ATAC-seq) are frequently used as a proxy for determining sites of TF binding within the genome (*15*). We therefore used ATAC-seq to profile regions of accessible chromatin in WT and KAP1 deletion ESCs. In agreement with the observed reduction of H3K9me3 heterochromatin and increase in H3K27ac at ERVs, we find that these regions become more accessible in KAP1 deletion ESCs (Fig. 2A). Loss of KAP1 resulted in an expansion of accessible regions specifically at IAP and RLTR10 elements (fig. S2A), active elements in the mouse genome previously shown to be silenced by KAP1 (*2, 16*–*18*). Inspection of two genes, *Pou5f1* and *Klf4*, showed that the regulatory elements encoding these genes became less accessible in the absence of KAP1 (Fig. 2B). Genome-wide, we found that H3K27ac-enriched regions showed a global loss of accessibility after KAP1 deletion (Fig. 2C). Interestingly, the gain in accessibility at H3K9me3-enriched regions appeared bimodal, in agreement with previous results that a subset of heterochromatin in ESCs is maintained by KAP1 in ESCs (Fig. 2C) (*2, 3*).

**Figure 2.**
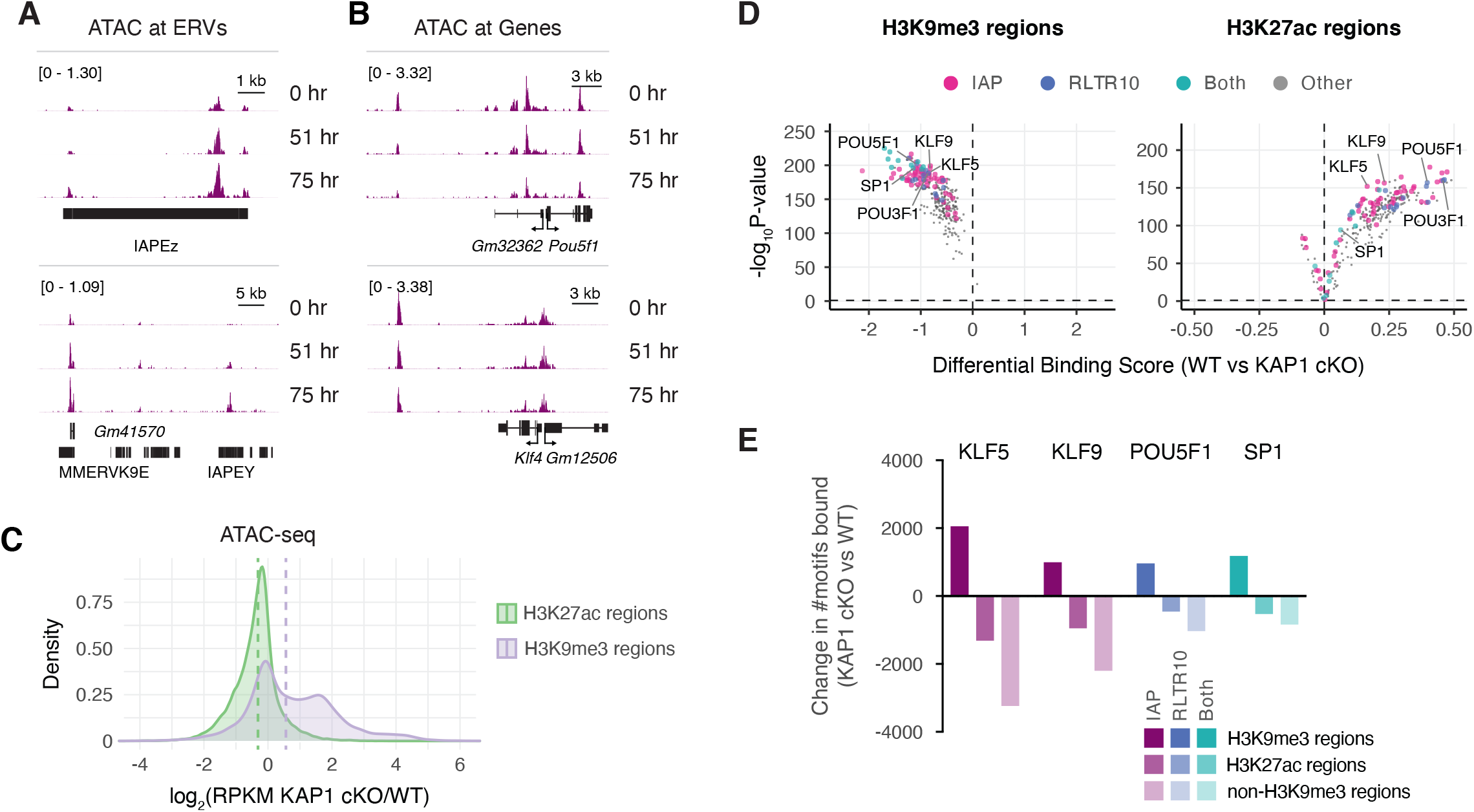
Heterochromatin at ERVs protects euchromatic chromatin accessibility. **(A)** Genome browser representations of ATAC-seq enrichment at ERVs in KAP1 cKO ESCs untreated (0 hr) or treated with 1 μM 4-OHT for 51 or 75 hrs. The y axis represents read density in counts per million (CPM). **(B)** Genome browser representations of ATAC-seq enrichment at genic regions in KAP1 cKO ESCs untreated (0 hr) or treated with 1 μM 4-OHT for 51 or 75 hrs. The y axis represents read density in counts per million (CPM). **(C)** Ratio (log2) of ATAC-seq enrichment at H3K9me3-enriched and H3K27ac-enriched regions in WT and KAP1 cKO ESCs. x axis values <0 indicate reduced enrichment in the absence of KAP1. **(D)** Pairwise comparison of TF activity at H3K9me3-enriched regions (left) or H3K27ac-enriched regions (right) between KAP1 cKO ESCs that were untreated or treated with 1 μM 4-OHT for 75 hrs. The volcano plot shows differential binding activity against the -log10(p value) for all investigated TF motifs. Each TF is represented by a single circle (n = 288). TF motifs enriched in KAP1 cKO ESCs have negative differential binding scores and TF motifs enriched in control ESCs have positive differential binding scores. Motifs enriched in specific repeat element families are color-coded as indicated. **(E)** Change in the number of bound motifs identified for indicated TFs at H3K9me3-enriched regions, H3K27ac-enriched regions, or non-H3K9me3-enriched regions in KAP1 cKO ESCs compared to WT ESCs. Positive values indicate a gain of TF binding at the indicated region in KAP1 cKO ESCs and negative values indicate a loss of TF binding.

In addition to measuring overall chromatin accessibility, ATAC-seq also contains regions of depleted signal within open chromatin that are protected from transposition by TF binding, referred to as TF footprints. We used a recently developed tool called TOBIAS (*19*) to perform comparative footprinting analysis of 288 expressed TFs with known consensus motifs in WT ESCs. This analysis confirmed that that specific repeat families are enriched with unique transcription factor binding motifs that become more bound upon KAP1 deletion (e.g., KLF family motifs are enriched within IAPs, POU family motifs are enriched within RLTR10s, and SP family motifs are enriched within both repeat classes) (fig. S2B) (*20, 21*). We then binned the genome into three groups: (1) regions enriched with H3K9me3 in WT ESCs and ATAC-seq signal in either WT or KAP1 deletion ESCs (H3K9me3 regions), (2) regions enriched with H3K27ac in WT ESCs and ATAC-seq signal in either WT or KAP1 deletion ESCs (H3K27ac regions), and (3) regions with ATAC-seq signal in either WT or KAP1 deletion ESCs but not enriched with H3K9me3 in WT ESCs (non-H3K9me3 regions). Comparison between K27ac and non-H3K9me3 regions allowed more or less stringency in our assessment of changes in chromatin accessibility at euchromatic regions in the absence of KAP1. Strikingly, KAP1 deletion had dramatically different effects on TF footprinting in heterochromatin and euchromatin. Specifically, we found that nearly all expressed TFs showed increased binding to their motifs in heterochromatic regions and reduced binding at euchromatic regions in KAP1 deleted ESCs (Fig. 2D, fig. S2C,D). Analysis of ATAC-seq data from WT and KAP1 cKO ESCs centered on motifs for selected TFs enriched in specific repeat elements showed clearly increased accessibility at heterochromatic regions and reduced accessibility at euchromatic regions upon loss of KAP1 (fig. S2E). These results were mirrored by the same directional changes in H3K27ac enrichment (fig. S2F). Computational assessment of the number of TF motifs predicted to be bound in each genomic region confirmed that specific TFs (e.g., KLF5, KLF9, POU3F1, POU5F1, SP1) gain binding sites in heterochromatic regions and concomitantly lose binding sites in euchromatic regions in the absence of KAP1 (Fig. 2E, fig. S2G). SETDB1 deletion had similar global effects on chromatin accessibility (fig. S3), supporting our hypothesis that heterochromatin at ERVs promotes euchromatic TF binding homeostasis.

Since ATAC-seq serves as a proxy for TF binding, we wanted to confirm these results with genomic assessment of TF binding itself. We performed CUT&Tag for two TFs whose motifs showed differential accessibility at heterochromatin and euchromatin in KAP1 cKO, KLF5 and KLF9 (Fig. 3, fig. S4). Overwhelmingly, these data support our ATAC-seq results, in that KAP1 deletion results in enhanced TF binding at ERVs containing motifs for KLF5 or KLF9 (Fig. 3A, fig. S4A) and more generally in regions enriched with H3K9me3 heterochromatin (Fig. 3B,C and fig. S4B,C). Likewise, KAP1 deletion reduces TF binding at genes containing motifs for KLF5 or KLF9 (Fig. 3A, fig. S4A) and more generally in regions enriched with H3K27ac, a marker of euchromatin (Fig. 3B,C and fig. S4B,C). Taken together, these data support a model in which loss of heterochromatin at ERVs reveals additional TF binding sites, potentially numbering in the hundreds to thousands, that compete with sites at euchromatic regions (e.g., genes and regulatory elements) for TF binding.

**Figure 3.**
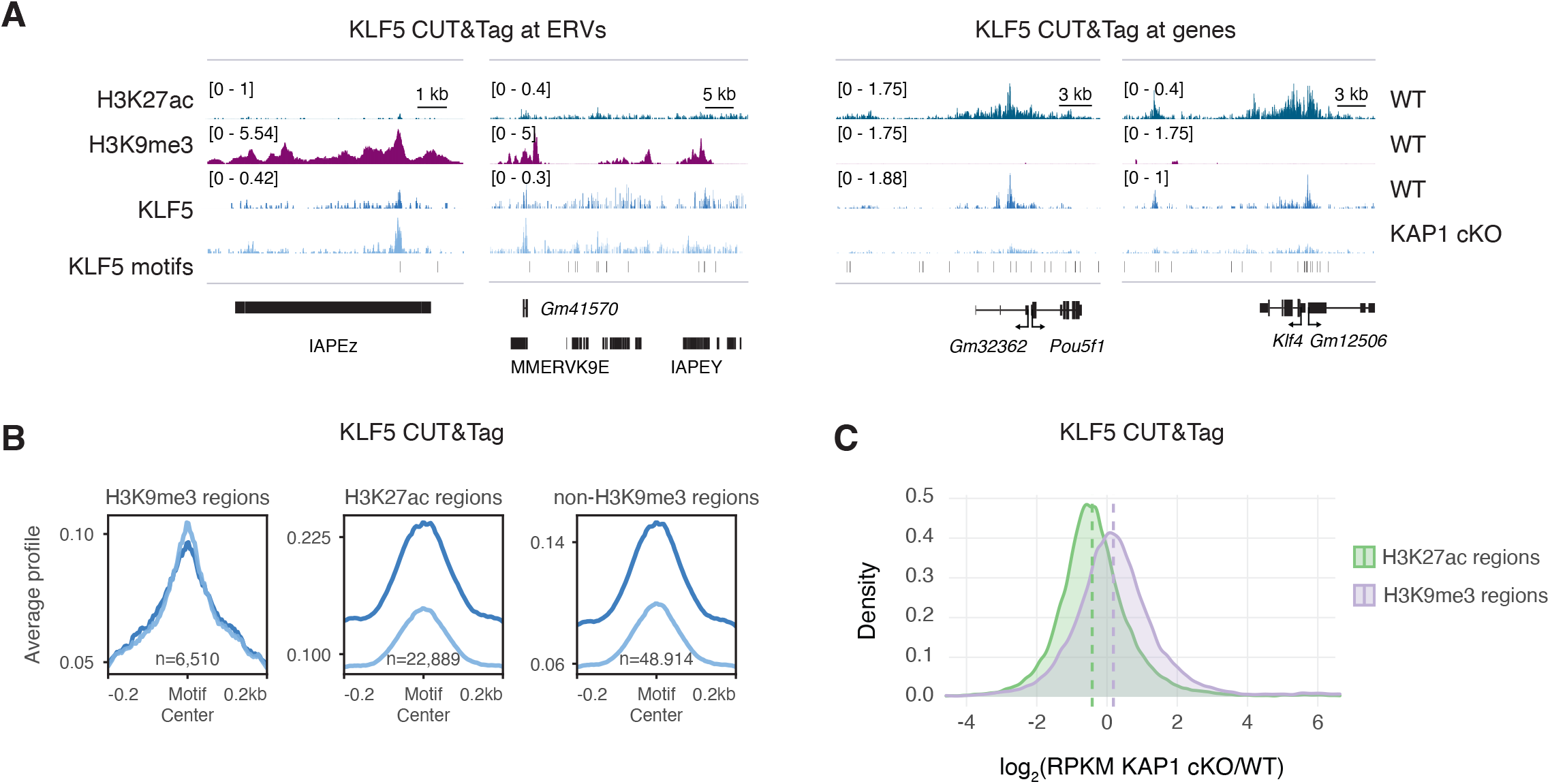
KAP1 deletion leads to reduced KLF5 binding at euchromatin. **(A)** Genome browser representations of KLF5 CUT&Tag enrichment at ERVs and genes in WT and KAP1 cKO ESCs. The y axis represents read density in counts per million (CPM). **(B)** KLF5 CUT&Tag average profiles at KLF5 motifs in H3K9me3-enriched regions (left), H3K27ac-enriched regions (center), and non-H3K9me3-enriched regions (right) in WT and KAP1 cKO ESCs. Data are centered on the motif and the number of motifs profiled are indicated. **(C)** Ratio (log2) of KLF5 CUT&Tag enrichment at H3K9me3-enriched and H3K27ac-enriched regions in WT and KAP1 cKO ESCs. x axis values <0 indicate reduced enrichment in the absence of KAP1.

Previous studies have reported that loss of KAP1 is lethal to developing embryos and that prolonged depletion causes loss of pluripotency in ESCs (*2, 3, 22, 23*). While ESCs acutely depleted of KAP1 maintained Oct4 protein levels over the course of our experiment (see Fig. 1), it is possible that changes in chromatin accessibility we observe are due to changes in the pluripotency regulatory landscape downstream of loss of stemness in KAP1 cKO ESCs. To precisely mimic the effects of exposing a large number of TF binding sites without the potentially confounding effects of KAP1 loss and global heterochromatin dysregulation, we designed a Piggybac transposon system to integrate exogenous binding site arrays (EBSAs) for a specific TF into ESCs (Fig. 4A). These arrays carry ∼150 copies of the motif for a single TF interspersed with 21 bp unique spacer sequences between each motif. As Piggybac is capable of inserting 10-40 copies of a transgene per cell, we predict that we can deliver upwards of 1500 copies or more of the TF motif to ESCs. Of note, this number is comparable to the number of individual motifs that become more bound in heterochromatin and less bound in euchromatin upon KAP1 deletion (see Fig. 2).

**Figure 4.**
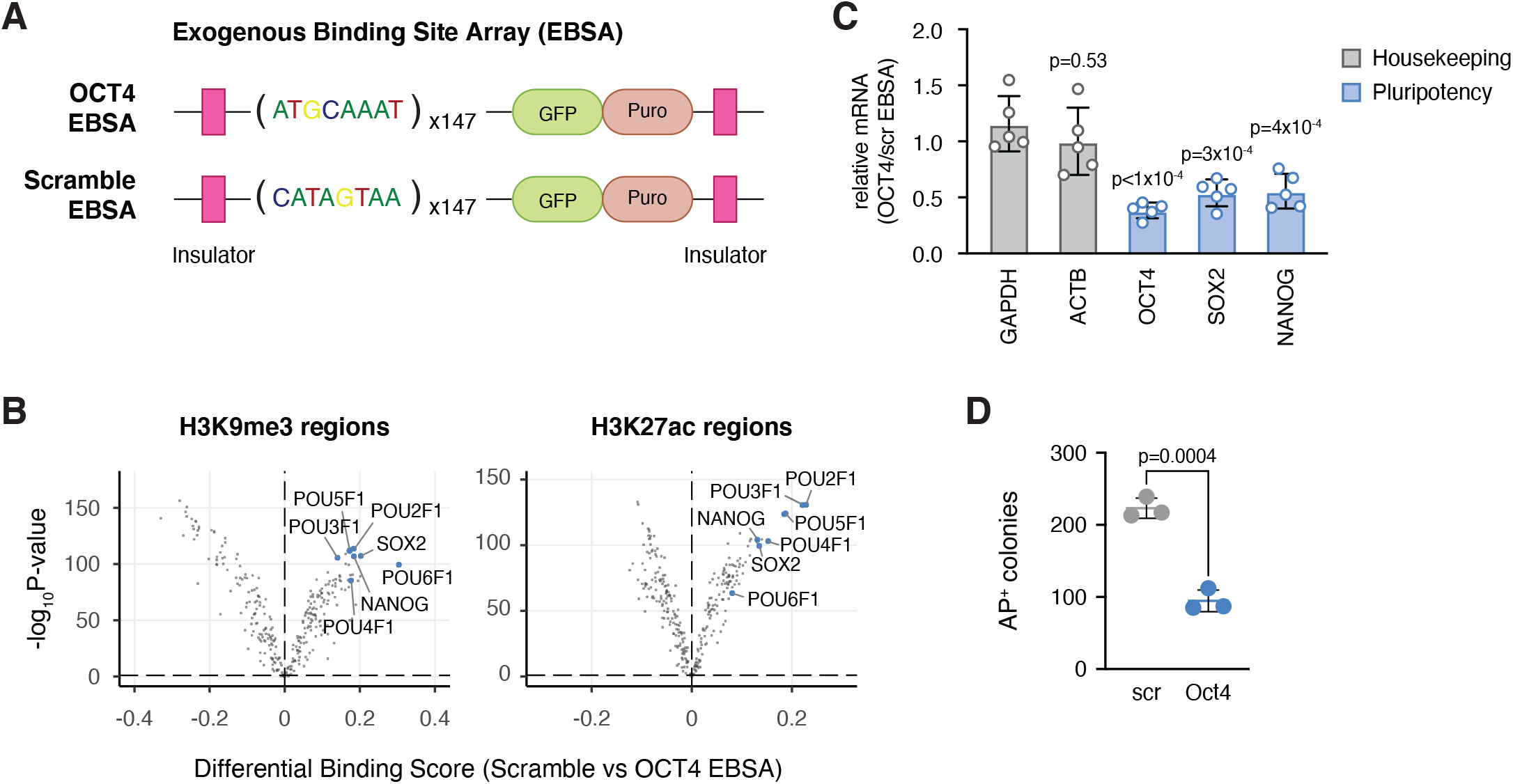
Exogenous TF binding site arrays mimic loss of ERV heterochromatin in ESCs. **(A)** Schematic of exogenous binding site arrays (EBSAs) containing 147 copies of either the POU5F1 (OCT4) TF motif or a scramble sequence. Motifs are interspersed with 21 bp spacer sequence. The Piggybac cassette expresses a GFP-T2A-Puro fusion and is flanked by insulator sequences. **(B)** Pairwise comparison of TF activity at H3K9me3-enriched regions (left) or H3K27ac-enriched regions (right) between ESCs transfected with scramble or OCT4 EBSAs. The volcano plot shows differential binding activity against the -log10(p value) for all investigated TF motifs. Each TF is represented by a single circle (n = 288). TF motifs enriched in OCT4 OBSA ESCs have negative differential binding scores and TF motifs enriched in scramble EBSA ESCs have positive differential binding scores. POU family motifs and select pluripotency motifs are labeled blue. **(C)** RT-qPCR analysis of housekeeping (grey) and pluripotency (blue) specific genes in OCT4 EBSA ESCs compared to scramble EBSA ESCs. Data represent mean ± s.d (*n* = 5). **(D)** Alkaline phosphatase positive (AP+) colonies formed from scramble and OCT4 EBSA ESCs seeded at clonal density.

We focused on the POU5F1 binding motif, as we predicted that redistribution of Oct4 TF binding away from euchromatic targets might not only be reflected by changes in chromatin accessibility but might also influence pluripotency, resulting in ESC differentiation (*24, 25*). We first performed a motif search of our 5 kb EBSA sequence to confirm the presence of the POU5F1 motif and the absence of any confounding TF motifs that may have been present in our spacer sequences (fig. S5A). Additionally, we confirmed that an array designed with a scrambled motif sequence (and identical spacer sequences) did not appreciably contain TF motifs that might affect interpretation of our experiment (e.g., the POU5F1-SOX2 motif was found 5 times in the control EBSA compared with 147 copies of the POU5F1 motif and 30 copies of the POU5F1-SOX2 motif in our designed EBSA) (fig. S5B). To determine how the EBSAs affect chromatin accessibility, we performed ATAC-seq from WT ESCs transfected with either the scramble EBSA or the OCT4 EBSA (Fig. 4B). Importantly, we found that chromatin accessibility at POU family motifs decreased in the OCT4 EBSA ESCs compared to scramble EBSA ESCs. In addition, these data show reduced chromatin accessibility at key pluripotency motifs, for example SOX2 and Nanog. Interestingly, POU5F1 TF footprinting was reduced in OCT4 EBSA ESCs at both euchromatic and heterochromatic regions, suggesting that the array is competing with OCT4 binding both at genic regulatory elements and at ERVs.

If the OCT4 EBSA is competing with OCT4 binding at its target genes, we might expect to observe reduced robustness of the transcriptional network directed by OCT4 (*26*). Accordingly, transfection with the OCT4 EBSA resulted in reduced expression of critical regulators of pluripotency (e.g., OCT4, SOX2, NANOG) without any appreciable effect on housekeeping genes such as beta actin or GAPDH (Fig. 4C). To further test the effect of competition for endogenous OCT4 protein on ESC pluripotency, we seeded OCT4 EBSA ESCs and scramble EBSA ESCs at clonal density and assessed their ability to form alkaline-phosphatase-positive (AP^+^) colonies, a marker of pluripotency. Strikingly, ESCs transfected with the OCT4 EBSA showed impaired capacity to form AP^+^ colonies compared with ESCs transfected with the scramble EBSA control (Fig. 4D, fig. 5SC).

## Discussion

Overall, our data support a model in which heterochromatin promotes TF binding within euchromatin by restricting TF access to ERVs. Of note, the genome undergoes two major waves of developmental epigenetic reprogramming (pre-implantation development and germ cell formation) during which ERVs become extensively activated in a highly programmed manner (*9, 27*). These time frames are also periods of dramatic transition in gene regulatory networks and transcription programs associated with a profound cellular plasticity required for lineage specification (*28*). We propose that one function of ERV activation during these times may be the presentation of competitive TF binding sites to weaken the robustness of existing transcription programs to allow the establishment of new programs, thus promoting transitions in cellular state.

There is precedent in the literature for viral competition for host factors. As with ERVs, latent viruses are rich with host-specific TF binding sites that drive transcription of the viral machinery. An emerging model termed “microcompetition” posits that these *cis*-regulatory elements within latent viruses compete with cellular genes for a limited amount of endogenous transcriptional machinery, leading to dysregulation of host transcription programs (*29*). Interestingly, a recent study found that copy number of latent EBV genome rather than infection correlated with viral oncogenicity, in support of this model (*30*). While studies of microcompetition are currently limited to infectious viruses, this model has striking implications for endogenous retroviral sequences, which make up roughly one tenth of mouse and human genomes.

Interestingly, previous studies have suggested that repetitive elements or even transgenes might serve as sinks for TF binding (*31*–*36*). Studies in bacteria have been particularly important in exploring the stoichiometry between TF protein level and TF binding sites and suggest that certain TFs may be more susceptible to titration effects than others (*35, 36*). While these studies provide important conceptual advances in our understanding of the biophysical determinants of transcription at the level of transcription factor binding, they use exogenous cellular perturbations and do not always mimic molecular mechanisms relevant to eukaryotic cells. Our study demonstrates that loss of heterochromatin at ERVs is required for TF redistribution. While it is possible that ERVs are competing with genes for transcription machinery such as the Mediator complex or RNAPII, the specificity of this competition must be driven by transcription factors themselves. Additionally, our model requires that occupied TF binding sites are at equilibrium with the molecules of TF available for binding (*35*–*37*). It is well-documented that not all predicted TF motifs in the genome are bound by TFs, and we still do not fully understand the “grammar” of regulatory elements that prefers binding at certain sites over others (*38*). Thus, a thorough molecular accounting of TF protein levels and improved predictive power of motif strength will no doubt improve future studies based on the model we put forth.

In closing, we propose that ERV derepression contributes to developmental transcriptional plasticity by competing with euchromatin for TF binding. Given the species-specific evolutionary arms race between host genomes and specific retroviral insertions, such a mechanism would require convergent evolution of this trait (*39*). Interestingly, heterochromatin dysregulation and loss of transcriptional homeostasis are both hallmarks of aging (*40*). Further, LINEs and ERVs are often hypomethylated and become derepressed in human cancer, where transcriptional plasticity is required for tumor formation and progression (*41*–*43*). Thus, our proposed mechanism may have broad implications not only for our understanding of developmental programs but also human health and disease.

## Acknowledgements

We thank members of the Banaszynski lab for helpful discussions; we thank D. Hancks and L. Kraus for helpful discussions and comments on this manuscript; UTSW BioHPC for computational infrastructure; UTSW McDermott Center for providing high-throughput sequencing services; UTSW Moody Foundation Flow Cytometry Core for FACS sorting services. L.A.B. is a Virginia Murchison Linthicum Scholar in Medical Research (UTSW Endowed Scholars Program) and an American Cancer Society Research Scholar.

## Author Contributions

R.O. and L.A.B. conceived and designed the study. R.O. performed experiments and computational analyses. L.A.B supervised and provided funding for the project.

R.O. and L.A.B prepared the figures and wrote the manuscript.

## Code Availability

Code to generate figures is available at https://github.com/utsw-medical-center-banaszynski-lab/xxx

## Data Availability

Datasets are deposited in the NCBI Gene Expression Omnibus using the following accession numbers: SuperSeries GSExx, ATAC-seq GSExxx, CUT&Tag GSExxx. Other datasets used for this study are available under GSE41903 (*4*), GSE151053 (*44*) and GSE114548 (*45*).

## Supplementary Materials

Materials and Methods

## Figure Legends

**Figure S1.**
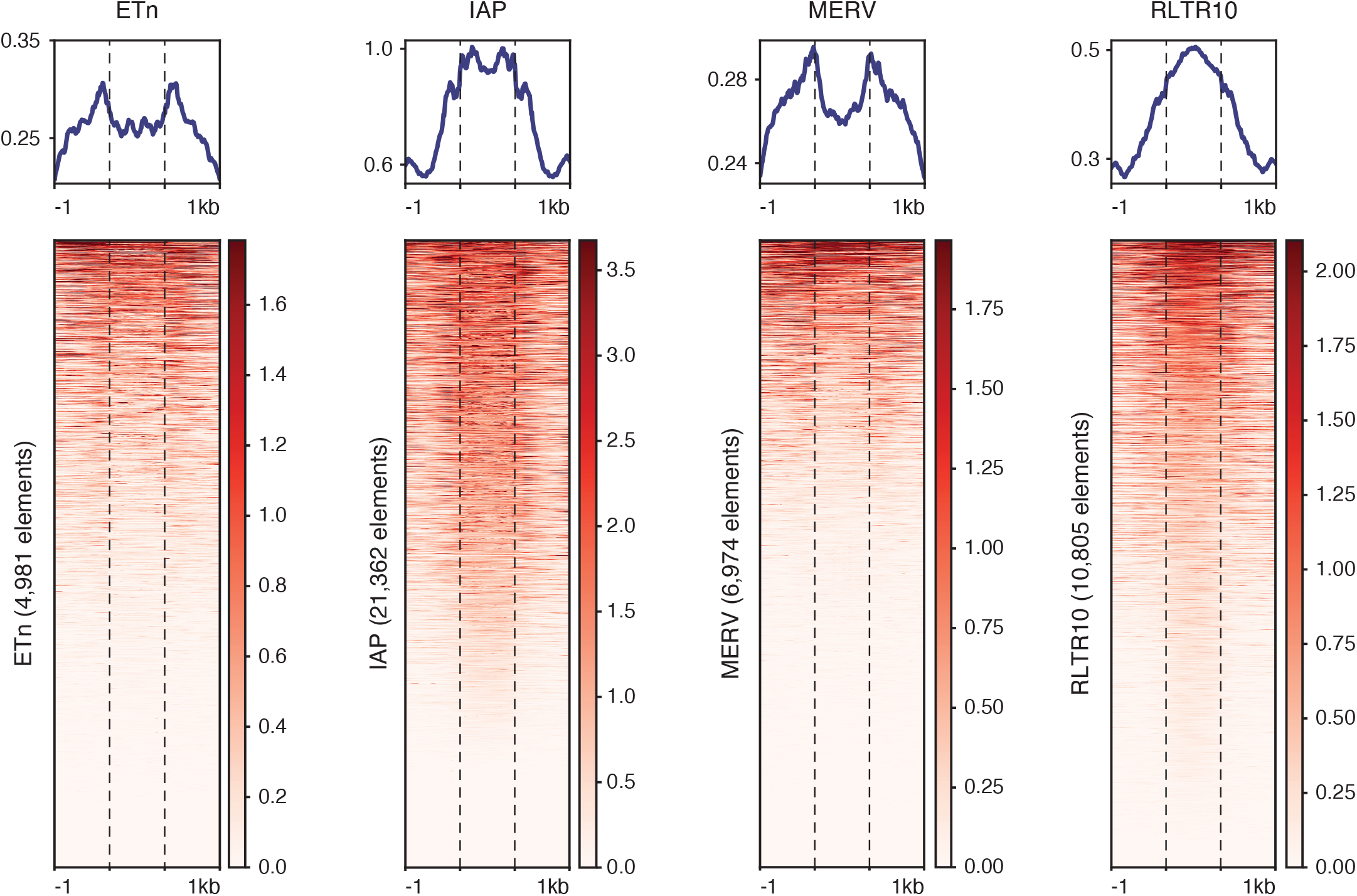
ERVs are enriched with H3K9me3. Related to Figure 1 Average profiles (top) and heatmaps (bottom) of H3K9me3 CUT&Tag at ETn, IAP, MERV, and RLTR10 endogenous retroelements in WT ESCs. ERVs are plotted as metagenes with 1 kb flanking sequence displayed for each analysis.

**Figure S2.**
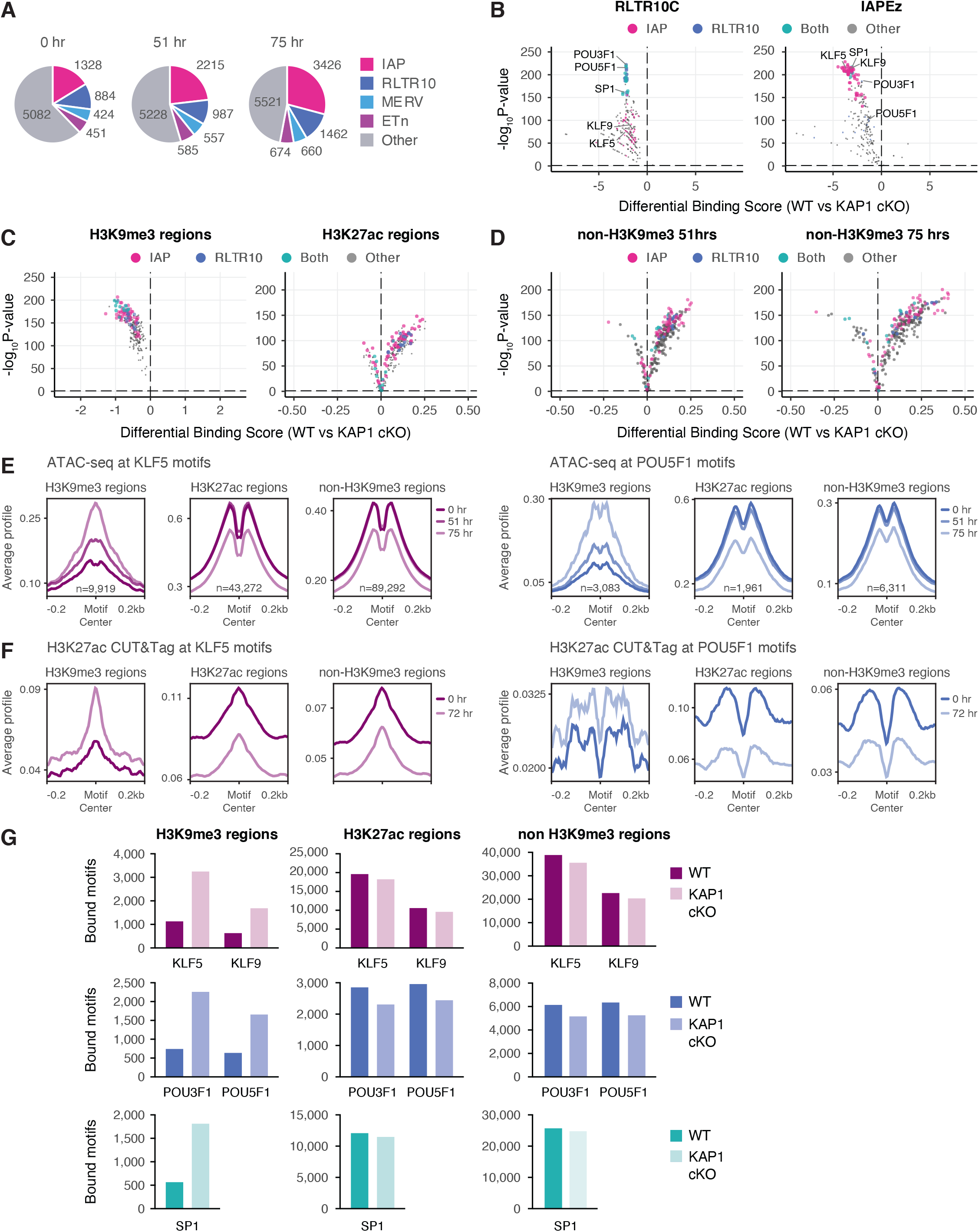
KAP1-mediated heterochromatin at ERVs protects euchromatic chromatin accessibility. Related to Figure 2 **(A)** Repeat element annotation of ATAC-seq regions of enrichment co-localizing with H3K9me3 enrichment in untreated control ESCs (left) or KAP1 cKO ESCs treated with 1 μM 4-OHT for 51 hr (center) or 75 hr (right). **(B)** Pairwise comparison of TF activity at RLTR10C elements (left) or IAPEz elements (right) between KAP1 cKO ESCs that were untreated or treated with 1 μM 4-OHT for 75 hrs. **(C)** Pairwise comparison of TF activity at H3K9me3-enriched regions (left) or H3K27ac-enriched regions (right) between KAP1 cKO ESCs that were untreated or treated with 1 μM 4-OHT for 51 hrs. **(D)** Pairwise comparison of TF activity at non-H3K9me3-enriched regions between KAP1 cKO ESCs that were untreated or treated with 1 μM 4-OHT for 51 hrs (left) or 75 hrs (right). For panels B-D, the volcano plot shows differential binding activity against the -log10(p value) for all investigated TF motifs. Each TF is represented by a single circle (n = 288). TF motifs enriched in KAP1 cKO ESCs have negative differential binding scores and TF motifs enriched in control ESCs have positive differential binding scores. Motifs enriched in specific repeat element families are color-coded as indicated. **(E, F)** ATAC-seq average profiles at **(E)** KLF5 and **(F)** POU5F1 motifs at H3K9me3-enriched regions (left), H3K27ac-enriched regions (center), and non-H3K9me3-enriched regions (right) in WT and KAP1 cKO ESCs. Data are centered on the motif and the number of motifs profiled are indicated. **(G, H)** H3K27ac CUT&Tag average profiles at **(G)** KLF5 and **(H)** POU5F1 motifs at H3K9me3-enriched regions (left), H3K27ac-enriched regions (center), and non-H3K9me3-enriched regions (right) in WT and KAP1 cKO ESCs. Data are centered on the motif and the number of motifs profiled are indicated in panels E and F. **(I)** Number of bound motifs identified for indicated TFs at H3K9me3-enriched regions, H3K27ac-enriched regions, and non-H3K9me3-enriched regions in KAP1 cKO ESCs compared to WT ESCs.

**Figure S3.**
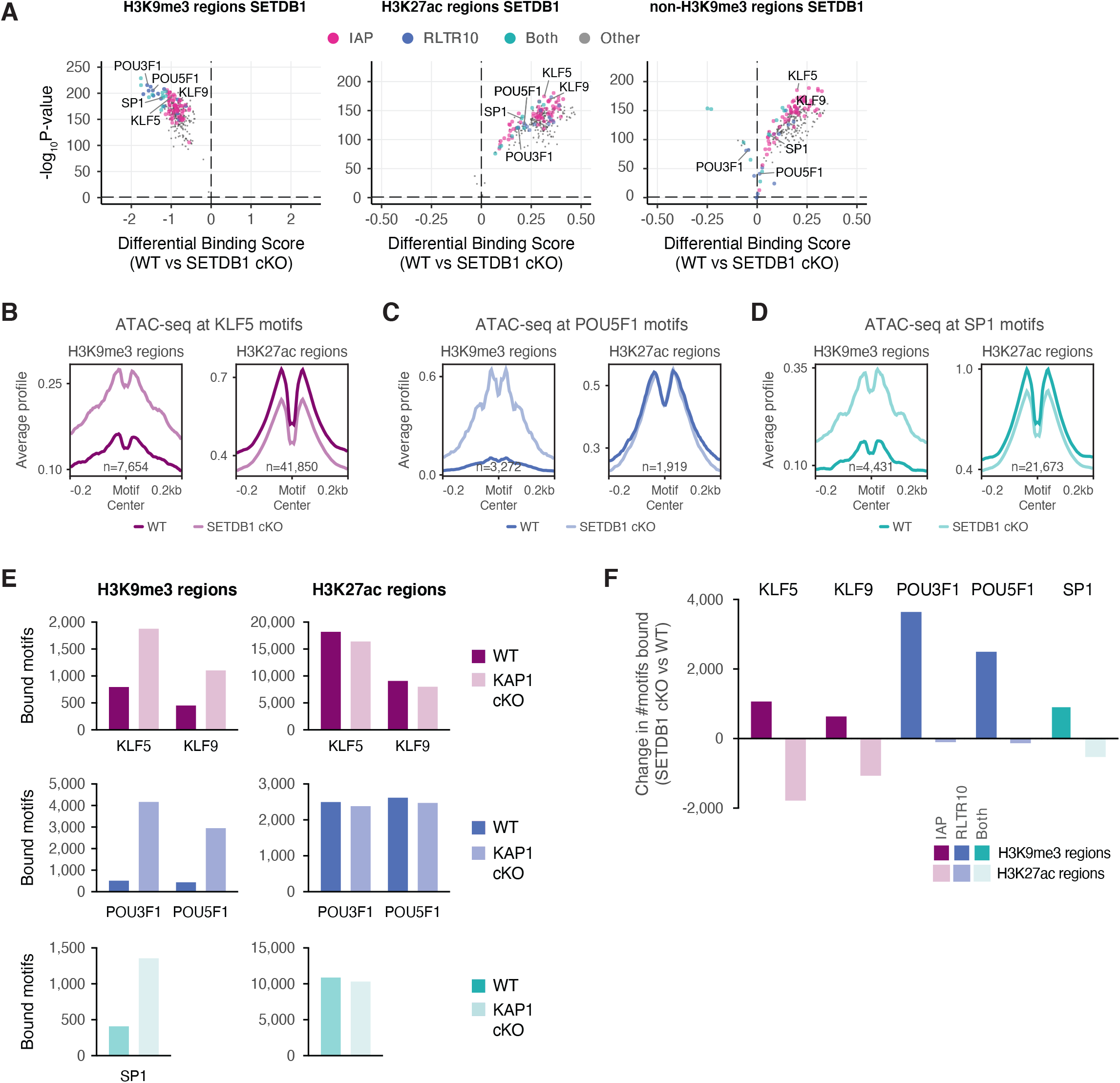
SETDB1-mediated heterochromatin at ERVs protects euchromatic chromatin accessibility. Related to Figure 2 **(A)** Pairwise comparison of TF activity at H3K9me3-enriched regions (left), H3K27ac-enriched regions (center), or non-H3K9me3-enriched regions (right) between SETDB1 cKO ESCs that were untreated or treated with 1 μM 4-OHT for 75 hrs. The volcano plot shows differential binding activity against the -log10(p value) for all investigated TF motifs. Each TF is represented by a single circle (n = 288). TF motifs enriched in SETDB1 cKO ESCs have negative differential binding scores and TF motifs enriched in control ESCs have positive differential binding scores. Motifs enriched in specific repeat element families are color-coded as indicated. **(B-D)** ATAC-seq average profiles at **(B)** KLF5, **(C)** POU5F1, and **(D)** SP1 motifs at H3K9me3-enriched regions (left) or H3K27ac-enriched regions (right) in WT and SETDB1 cKO ESCs. Data are centered on the motif and the number of motifs profiled are indicated. **(E)** Number of bound motifs identified for indicated TFs at H3K9me3-enriched regions and H3K27ac-enriched regions in SETDB1 cKO ESCs compared to WT ESCs. **(F)** Change in the number of bound motifs identified for indicated TFs at H3K9me3-enriched regions and H3K27ac-enriched regions in SETDB1 cKO ESCs compared to WT ESCs. Positive values indicate a gain of TF binding at the indicated region in SETDB1 cKO ESCs and negative values indicate a loss of TF binding.

**Figure S4.**
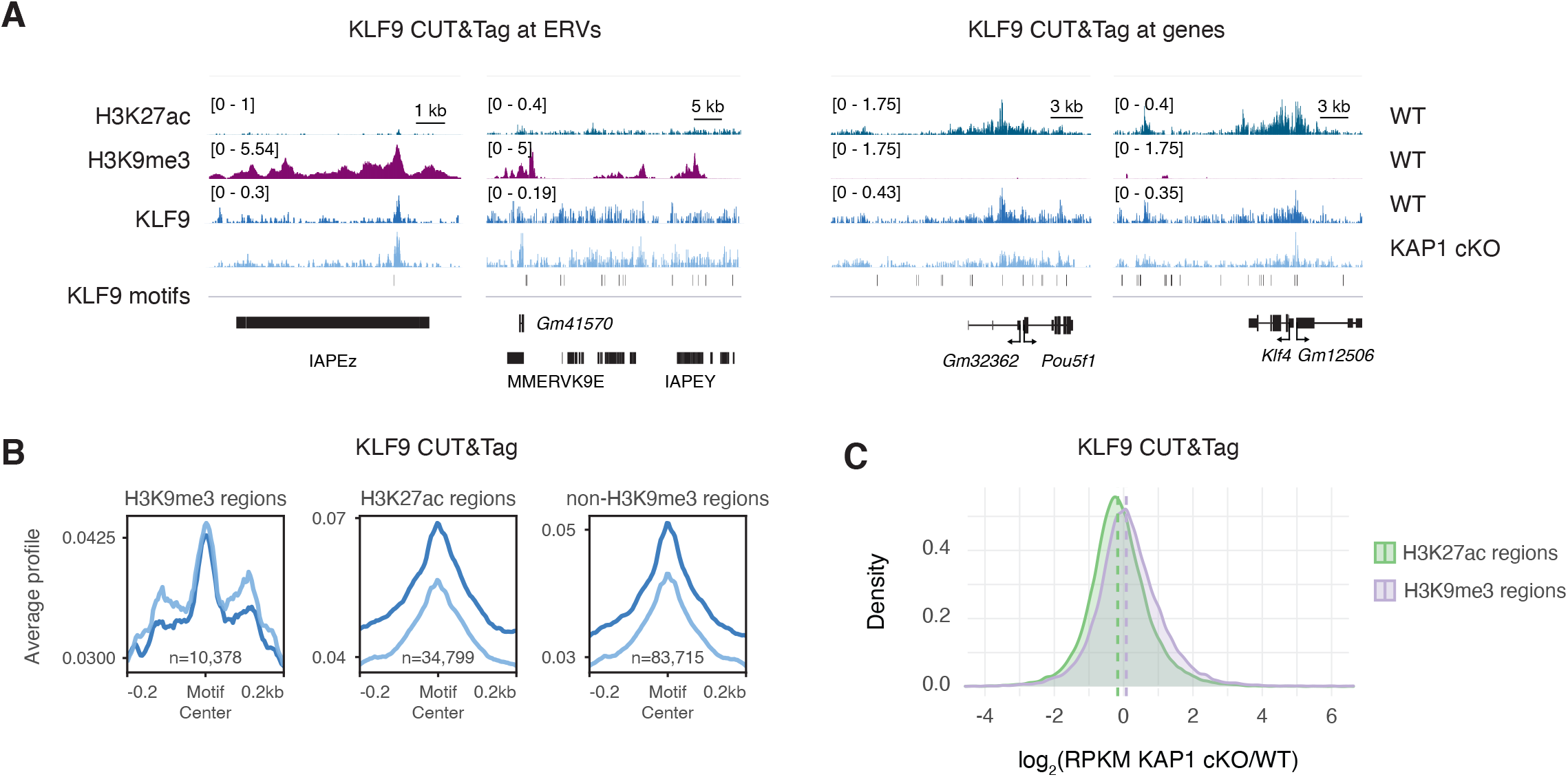
KAP1 deletion leads to reduced KLF9 binding at euchromatin. Related to Figure 3 **(A)** Genome browser representations of KLF9 CUT&Tag enrichment at ERVs and genes in WT and KAP1 cKO ESCs. The y axis represents read density in counts per million (CPM). **(B)** KLF9 CUT&Tag average profiles at KLF9 motifs in H3K9me3-enriched regions (left), H3K27ac-enriched regions (center), and non-H3K9me3-enriched regions (right) in WT and KAP1 cKO ESCs. Data are centered on the motif and the number of motifs profiled are indicated. **(C)** Ratio (log2) of KLF9 CUT&Tag enrichment at H3K9me3-enriched and H3K27ac-enriched regions in WT and KAP1 cKO ESCs. x axis values <0 indicate reduced enrichment in the absence of KAP1.

**Figure S5.**
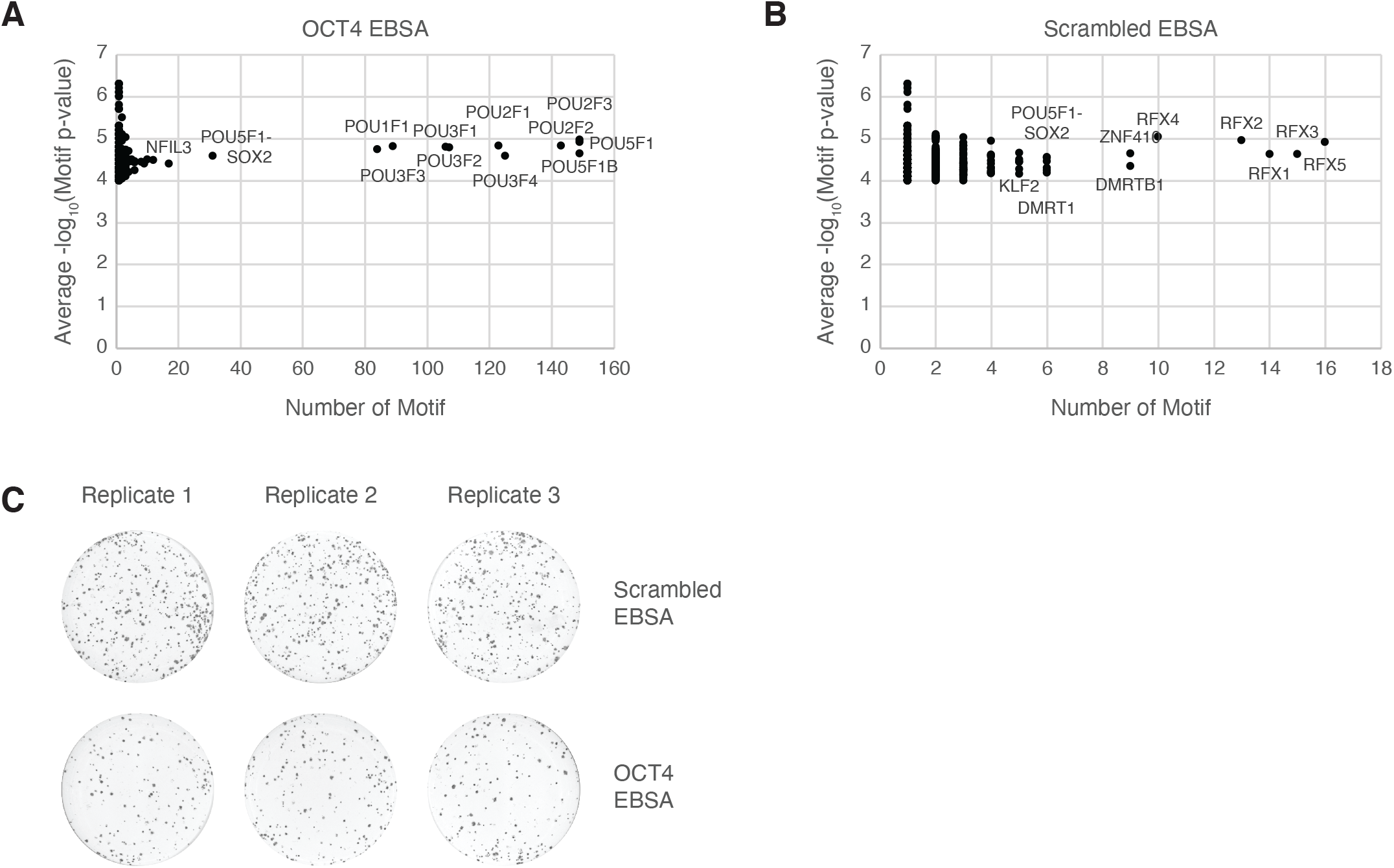
Exogenous binding site arrays carrying the OCT4 motif affect ESC pluripotency. Related to Figure 4 **(A)** Homer prediction of TF motifs present in the OCT4 exogenous binding site array (EBSA). **(B)** Homer prediction of TF motifs present in the scramble EBSA. **(C)** Alkaline phosphatase staining of scramble and OCT4 EBSA ESCs seeded at clonal density.

## Materials and Methods

### ESC culture

ESCs were maintained under standard conditions on gelatin-coated plates at 37 °C, 5% CO_2_, and ambient O_2_ in Knockout DMEM (Thermo Fisher) supplemented with NEAA, GlutaMAX, penicillin/streptomycin (Thermo Fisher), 15% ESC-screened fetal bovine serum (Hyclone), 0.1 mM 2-mercaptoethanol (Fisher), and leukemia-inhibitory factor (LIF). Generation of KAP1 cKO and ESET cKO ESCs has been described previously (*2, 3*). KAP1 or ESET deletion was induced with 1 μM 4-OHT (Sigma) treatment for two days in cKO ESCs. ESCs were routinely screened for mycoplasma.

### Antibodies

KAP1 (ab10483, Abcam), OCT4 (sc-5279, Santa Cruz), H3 general (ab1791, Abcam), anti-mouse IgG-HRP (NA93V, GE, Lot # 9773218), anti-rabbit IgG-HRP (170-6515, Biorad, Lot # 350003248), H3K9me3 (ab8898, Abcam), H3K27ac (39133, Active Motif), KLF5 (61099, Active Motif), KLF9 (701888, Thermo Fisher).

### Western Blotting

ESCs were lysed with micrococcal nuclease (Worthington) in 50 mM Tris pH 7.5, 1 mM CaCl2, 0.2% Triton X-100 and 5 mM sodium butyrate. Proteins from whole cell lysate were separated in Laemmli buffer by SDS-PAGE and transferred to PVDF membranes (Millipore). Membranes were blocked in 5% milk in Tris-buffered saline with 0.1% Tween-20 (TBST) and incubated with primary antibodies overnight at 4 °C or for 2 hours at room temperature. Membranes were washed with TBST, incubated with HRP-conjugated secondary antibodies for 1 h, incubated with HRP substrate (Fisher) and imaged using a ChemiDoc MP Imaging System (BioRad).

### Chromatin Immunoprecipitation (ChIP)

#### Native ChIP

10^7^ cells were trypsinized, washed, then lysed (50 mM TrisHCl pH 7.4, 1 mM CaCl2, 0.2% Triton X-100, 10 mM NaButyrate, and protease inhibitor cocktail (Roche)) with micrococcal nuclease (Worthington) for 5 min at 37 °C to recover mono-to tri-nucleosomes. Nuclei were lysed by brief sonication and dialyzed twice into RIPA buffer (10 mM Tris pH 7.6, 1 mM EDTA, 0.1% SDS, 0.1% Na-Deoxycholate, 1% Triton X-100) for 1 hr at 4 °C. 5% of soluble material was reserved as input DNA. 5 μg of antibody bound to 50 μl protein A or protein G Dynabeads (Invitrogen) was incubated with soluble chromatin overnight at 4 °C. Magnetic beads were washed as follows: 3x RIPA buffer, 2x RIPA buffer + 300 mM NaCl, 2x LiCl buffer (250 mM LiCl, 0.5% NP-40, 0.5% NaDeoxycholate), 1x TE + 50 mM NaCl. Chromatin was eluted and treated with RNaseA and Proteinase K. ChIP DNA was purified using QIAquick PCR Purification Kit (Qiagen).

#### ChIP-qPCR

PCR was performed in triplicate using a LightCycler® 480 Instrument II system and Power SYBR Green PCR master mix. ChIP DNA samples were diluted 1:50 in H2O, with 5 μl used per reaction. ChIP-qPCR signal is represented as percent input.

The following primer sequences were used in this study:

IAPA

F: GCGCAAGGAAGATCCCTCAT

R: TTATTACCGCCGTTCCCCAG

IAPEz

F: CTAGGAGATGCTCGTCGCTG

R: AATGGGCTGCTTCTTCCTCC

eOCT4

F: CTCTCGTCCTAGCCCTTCCT

R: CATGGGTCCAAGTCATCCCC

eKLF4

F: AGGTGAGCGGTGGGTATAGA

R: CCCCAGATTCCAGGCTTTGT

#### Quantitative RT-PCR

mRNA was isolated using QIAGEN RNeasy. 500 ng of total RNA was reverse transcribed using random hexamers and MultiScribe reverse transcriptase. mRNA expression was analyzed by quantitative PCR (qPCR) with SYBR Green using a LightCycler 480 (Roche).

The following primer sequences were used in this study:

ACTB

F:AACCCTAAGGCCAACCGTGAAAAG

R:TGGCGTGAGGGAGAGCATAGC

GAPDH

F:GGTTGTCTCCTGCGACTTCAACAGC

R:CGAGTTGGGATAGGGCCTCTCTTGC

NANOG

F: TTGCTTACAAGGGTCTGCTACT

R: ACTGGTAGAAGAATCAGGGCT

POU5F1 (OCT4)

F: TTGGGCTAGAGAAGGATGTGGTT

R: GGAAAAGGGACTGAGTAGAGTGTGG

SOX2

F: CGCGGCGGAAAACCAAGACG

R: GCCGGCGCCCACCCCAACC

#### ATAC-Seq

The modified ATAC-sequencing protocol, Omni-ATAC was performed as previously described (*46*). Briefly, 10^5^ cells were lysed with resuspension buffer (Tris 10 mM, pH 7.4, 10 mM NaCl, 3 mM MgCl2, 0.1% NP-40, 0.1% Tween-20, and 0.01% Digitonin) and nuclei were collected for tagmentation at 37 °C for 30 minutes (Illumina Tagment DNA Enzyme and Buffer Small Kit). The reaction was immediately purified using Qiaquick PCR Purification Kit (Qiagen) and eluted in 20 μl water. Eluted DNA was amplified using NEBNext Ultra II PCR Master Mix (NEB) and purified using AMPure XP beads. Samples were pooled for multiplexing and sequenced using paired-end sequencing on the Illumina NextSeq 500.

#### Genomics analysis

Quality of ChIP-seq datasets was assessed using the FastQC tool. ChIP-seq raw reads were adapter and quality trimmed using Trimgalore (*47*). Trimmed reads were aligned to the mouse reference genome (mm10) with Bowtie2 (*48*) (bowtie2 -q -R 3 -N 1 -L 20 -i S,1,0.50 --end-to-end --dovetail --no-mixed -X 2000). Multimapping reads were randomly assigned. Optical duplicate reads were filtered using Picard. Reads which mapped to the mitochondrial genome were removed with Samtools (*49*) (samtools idxstats $sample.sorted.bam | cut -f 1 | grep -v chrM | xargs samtools view -b $sample.sorted.bam). Peak calling was performed with MACS2 software (*50*) (--keep-dup all --nomodel -B -f BAMPE, --broad peakcalling was used for H3K9me3 and H3K27ac ChIPs) and an FDR cutoff of 0.001 was applied to generate peak bedfiles. Peaks which intersected blacklisted high-signal genomic regions were removed. BigWig files were generated from alignments using deepTools (*51*) and normalized to counts per million (CPM). Visualization of bigWigs was done in Integrative Genomics Viewer (*52*). Intersections between different peak sets were made using BEDTools (*53, 54*). Heatmaps and average profiles were generated using deepTools. Density plots representing read densities and fold-changes in read densities were generated using R and ggplot2 (H. Wickham. ggplot2: Elegant Graphics for Data Analysis. Springer-Verlag New York, 2016).

#### H3K9me3, H3K27ac, and non-H3K9me3 regions

A merged peak file was generated for KAP1 and SETDB1 (GSE151053) (*55*) containing all WT and KO peaks in the respective condition. H3K9me3 regions were generated by intersecting merged ATAC peaks (either KAP1 or SETDB1) with H3K9me3 Cut and Tag peaks (in WT ESCs). H3k27ac regions were identified by intersecting the remaining ATAC peaks with peaks called from previously published H327ac ChIP data (GSE114548) (*45*). Non-H3K9me3 regions were generated by removing the K9me3 peak set from respective KAP1 and SETDB1 ATAC peaks.

#### Motif analysis

Previously published KAP1 cKO RNA-seq data was analyzed using DESeq2 (*56*). Expressed transcription factors were identified from RNA-seq (GSE41903) (*4*) using a baseMean cut-off of >74 (the median expression of all identified TFs). Position frequency matrices for expressed TF motifs were downloaded from JASPAR database (*57*). Motif bedfiles (KLF5 and KLF9) were generated using FIMO (*58*) with a *p* value cutoff of 0.0002.

#### TOBIAS analysis

We generated merged peak files containing all ATAC peaks present in each condition of every TOBIAS comparison. Replicate bam files were merged using Samtools. TOBIAS pipeline (*19*) was used to generate bigWig files scored for TF binding. BINDetect was then used to generate pairwise differential TF binding scores between merged samples for each expressed TF motif. For analysis of differential binding scores specifically in H3K9me3, H3K27ac, or non-H3K27ac regions, BINDetect was restricted using option --output-peaks to the indicated peaks.

#### Transfection of EBSAs

4 μg OCT4 or scramble EBSA vector and 1 μg vector containing Piggybac transposase were transfected in Opti-MEM media into 4×10^6^ WT ESCs using Lipofectamine 3000. At 48 hours post-transfection, cells were either sorted for FACS analysis or replated with 1 μg/ml puromycin. Sorted GFP-High cells were used for subsequent ATAC-seq and qPCR analyses.

#### Colony Formation Assay

Four days post-puromycin treatment, ESCs transfected with either OCT4 or scramble motif EBSAs were seeded at a density of 50K cells/well in 6 well-plates. Media was changed every 2 days. Five days after initial seeding, cells were fixed and stained using the Alkaline Phosphatase Staining Kit (STEMGENT, 00–0055). The colonies were quantified using ImageJ’s particle function analysis and technical triplicates were averaged for each condition.

